# Local homeostatic regulation of the spectral radius of echo-state networks

**DOI:** 10.1101/2020.07.21.213660

**Authors:** Fabian Schubert, Claudius Gros

**Affiliations:** Institute for Theoretical Physics, Goethe University Frankfurt, Frankfurt am Main, Hesse, Germany

## Abstract

Recurrent cortical network dynamics plays a crucial role for sequential information processing in the brain. While the theoretical framework of reservoir computing provides a conceptual basis for the understanding of recurrent neural computation, it often requires manual adjustments of global network parameters, in particular of the spectral radius of the recurrent synaptic weight matrix. Being a mathematical and relatively complex quantity, the spectral radius is not readily accessible to biological neural networks, which generally adhere to the principle that information about the network state should either be encoded in local intrinsic dynamical quantities (e.g. membrane potentials), or transmitted via synaptic connectivity. We present two synaptic scaling rules for echo state networks that solely rely on locally accessible variables. Both rules work online, in the presence of a continuous stream of input signals. The first rule, termed flow control, is based on a local comparison between the mean squared recurrent membrane potential and the mean squared activity of the neuron itself. It is derived from a global scaling condition on the dynamic flow of neural activities and requires the separability of external and recurrent input currents. We gained further insight into the adaptation dynamics of flow control by using a mean field approximation on the variances of neural activities that allowed us to describe the interplay between network activity and adaptation as a two-dimensional dynamical system. The second rule that we considered, variance control, directly regulates the variance of neural activities by locally scaling the recurrent synaptic weights. The target set point of this homeostatic mechanism is dynamically determined as a function of the variance of the locally measured external input. This functional relation was derived from the same mean-field approach that was used to describe the approximate dynamics of flow control.

The effectiveness of the presented mechanisms was tested numerically using different external input protocols. The network performance after adaptation was evaluated by training the network to perform a time delayed XOR operation on binary sequences. As our main result, we found that flow control can reliably regulate the spectral radius under different input statistics, but precise tuning is negatively affected by interneural correlations. Furthermore, flow control showed a consistent task performance over a wide range of input strengths/variances. Variance control, on the other side, did not yield the desired spectral radii with the same precision. Moreover, task performance was less consistent across different input strengths.

Given the better performance and simpler mathematical form of flow control, we concluded that a local control of the spectral radius via an *implicit* adaptation scheme is a realistic alternative to approaches using classical “set point” homeostatic feedback controls of neural firing.

**Author summary:** How can a neural network control its recurrent synaptic strengths such that network dynamics are optimal for sequential information processing? An important quantity in this respect, the spectral radius of the recurrent synaptic weight matrix, is a non-local quantity. Therefore, a direct calculation of the spectral radius is not feasible for biological networks. However, we show that there exist a local and biologically plausible adaptation mechanism, flow control, which allows to control the recurrent weight spectral radius while the network is operating under the influence of external inputs. Flow control is based on a theorem of random matrix theory, which is applicable if inter-synaptic correlations are weak. We apply the new adaption rule to echo-state networks having the task to perform a time-delayed XOR operation on random binary input sequences. We find that flow-controlled networks can adapt to a wide range of input strengths while retaining essentially constant task performance.

## Introduction

Echo state networks are a class of recurrent neural networks that can be easily trained on time dependent signal processing tasks. Once externally induced, the recurrent activity provides a reservoir of non-linear transformations [1], both in time and space, that may be utilized by a linear readout unit. Training the linear output units makes an echo state network a highly effective prediction machine [1, 2]. However, achieving optimal performance needs fine tuning of the network properties, in particular the spectral radius *R* = |Λ_*max*_|. It is closely related to the so-called echo state property [3]. While different equivalent formulations exist, a common definition for the echo-state property posits that specific input sequences lead to uniquely determined sequences of neural activity. This presumption implies that perturbations of the network’s initial conditions decay over time. Theoretical approaches linking the echo-state property to network parameters have led to results that can be divided into necessary and sufficient conditions for obtaining the echo state property. In particular, early works on echo state networks already established coarse bounds for the choice of the recurrent weight matrix, namely *R* < 1 as a necessary condition and *σ*_max_ < 1 as a sufficient condition, where *σ*_max_ is the largest singular value [3]. In practice, setting *R* ≈ 1 has proven to be sufficient for achieving the echo state property and good network performance [1, 3, 4]. This observation was supported by a theoretical result stating that for *R* < 1, the probability for the network dynamics being contracting tends to 1 for a large number of nodes [5]. If, however, *R* is significantly smaller than 1, information is lost too fast over time, which is detrimental for tasks involving sequential memory. A spectral radius of about one is hence best [6], in the sense that it provides a maximal memory capacity if the network operates in a linear regime [7, 8]. Similarly, Boedecker et al. found the best memory capacity to be close to the largest Lyapunov exponent being equal to zero [9], which can be understood as a generalization of the aforementioned results for nonlinear dynamics.

Aside their applications as efficient machine learning algorithms, echo state networks are potentially relevant as models of information processing in the brain [10–12].

An extension to layered ESN architectures was presented by Gallicchio and Micheli [13], which bears greater resemblance to the hierarchical structure of cortical networks than the usual shallow ESN architecture. This line of research illustrates the importance to examine whether local and biological plausible principles exist that would allow to tune the properties of the neural reservoir to the “edge of chaos” [14], in particular when a continuous stream of inputs is present. The rule has to be independent of the network topology, which is not a locally accessible information, and of the distribution of synaptic weights.

Generally speaking, strategies for optimizing ESN hyperparameters can be divided in two categories: supervised and unsupervised methods, where the first one utilizes an error signal, while the latter uses only information contained within the network dynamics. Our work is concerned with unsupervised optimization, independent of the actual task at hand. Several studies investigated local unsupervised intrinsic plasticity rules for improving neural reservoirs [15–17], usually by defining a target output distribution that each neuron attempts to reproduce by changing neural gains and biases, or a target distribution for the local optimization of information transmission [18, 19]. In general, however, it is difficult to formulate an explicit analytic theory that quantifies the relation between intrinsic neural parameters and global network dynamics, or corresponding parameters, as, for example, the spectral radius. One issue with choosing a fixed target distribution for optimizing network performance is that changing the external input, e.g. its mean or variance, affects the configuration of gains and biases resulting from the intrinsic plasticity mechanism. It can happen that the induced changes decrease performance by driving the network away from the critical point, which is known to be a beneficial dynamical state for sequence learning tasks [20–25].

Here, we propose and compare two unsupervised homeostatic mechanisms, which we termed flow control and variance control. Both are supposed to regulate the mean and variance of neuronal firing such that the network works in an optimal regime with regard to sequence learning tasks. The mechanisms act on two sets of node-specific parameters, the biases *b*_*i*_ and the neural gain factors *a*_*i*_. This approach can be considered a realization of dual homeostasis, which has been investigated previously with respect to a stable control of the mean and the variance of neural activity [26]. In this framework, the adaptation of the bias acts an intrinsic plasticity for the control of the internal excitability of a neuron [27–29], while the gain factors functionally correspond to a synaptic scaling of the recurrent weights [30–32].

We restricted ourselves to ‘biologically plausible’ adaption mechanisms, viz mechanisms for which the dynamics of all variables is local, i.e., bound to a specific neuron. Additional variables enter only when locally accessible. In a strict sense, this implies that local dynamics is determined exclusively by the dynamical variables of the neuron and by information about the activity of afferent neurons. Being less restrictive, one could claim that it should also be possible to access aggregate or ‘mean-field’ quantities that average a property of interest with respect to the population. For example, nitric oxide is a diffusive neurotransmitter that can act as a measure for the population average of neural firing rates [33].

In this study, we used standard discrete-time dynamics, which is markovian, with information about past states being stored dynamically. Following a general description of the model and the network, we present the flow control: it involves a normalization constraint in phase space (Sect. *Autonomous spectral radius regulation*). We evaluate the performance of networks that were subject to adaptation in Sect. *XOR-memory recall*, using a nonlinear recall task based on a sequential binary XOR-operation. Finally, we discuss the influence of node-to-node cross-correlations within the population in Sect. *Input induced correlations*.

## Model

A full description of the network model and parameters can be found in the methods section. We briefly introduce the network dynamics as

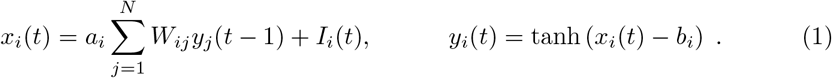

Each neuron’s membrane potential *x*_*i*_ consists of a recurrent contribution 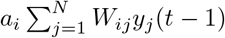 and some external input *I*_*i*_(*t*). The biases *b*_*i*_ were subject to the following homeostatic adaptation:

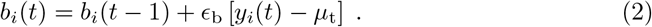

Here, *μ*_t_ defines a target for the average activity.

For *a*_*i*_, we considered two different forms of update rules. The first, flow control, is given by

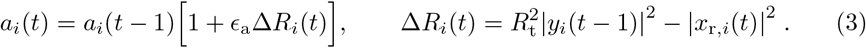

The parameter *R*_t_ defines the desired target spectral radius. We also considered an alternative global update rule where Δ*R*_*i*_(*t*) was given by

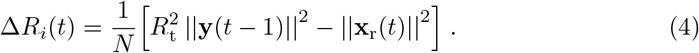

However, since this is a non-local rule, it only served as a comparative model to Eq. (3) when we investigated the effectiveness of the adaptation mechanism.

The second rule, variance control, has the form

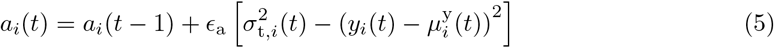

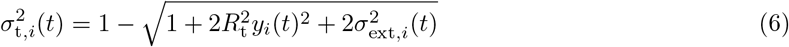

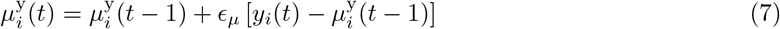

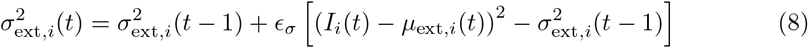

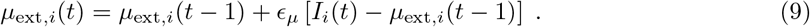

Eq. (5) drives the average variance of each neuron towards a desired target variance. While Eq. (7)–(9) are simply local trailing averages of the mean activity and the mean and variance of the external input, Eq. (6) is an analytic expression that was derived from a mean field approximation which is explained in S2 Appendix. Similar to flow control, we also considered a non-local version for comparative reasons where (6) was replaced with

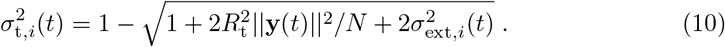

Before proceeding to the results, we explain the mathematical background of the proposed adaptation rules in more detail.

### Autonomous spectral radius regulation

We would like to explain the theoretical framework that led us to propose Eq. (3) and (6) as a regulatory mechanism for the spectral radius of the recurrent weight matrix.

The circular law of random matrix theory states that the eigenvalues *λ*_*j*_ are distributed uniformly on the complex unit circle if the elements of a real *N* × *N* matrix are drawn from distributions having zero mean and standard deviation 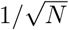 [34]. Noting that the internal weight matrix *W*_*ij*_ has *p*_r_*N* non-zero elements per row, the circular law implies that the spectral radius of *a*_*i*_*W*_*ij*_, the maximum of |*λ*_*j*_|, is unity when the synaptic scaling factors *a*_*i*_ are set uniformly to 1*/σ*_w_. Our goal is to investigate adaption rules for the synaptic rescaling factors that are based on dynamic quantities, which includes the membrane potential *x*_*i*_, the neural activity *y*_*i*_ and the input *I*_*i*_.

A *N* × *N* matrix with i.i.d. entries with zero mean and 1*/N* variance will have a spectral radius of one as *N* → ∞. Rajan and Abbott [35] investigated the case where the statistics of the columns of the matrix differ in their means and variances: given row-wise E-I balance for the recurrent weights, the square of the spectral radius of a random *N* × *N* matrix whose columns have variances 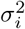 is 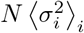 for *N* → ∞. Since the eigenvalues are invariant under transposition, this result can also be applied to our case, where node-specific gain factors *a*_*i*_ were applied to each *row* of the recurrent weights. Thus, the spectral radius *R*_a_ of random matrices *a*_*i*_*W*_*ij*_ is approximately given by

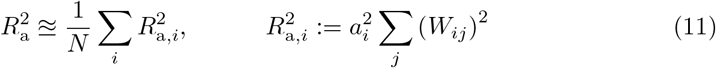

for large *N*, assuming that the distribution underlying 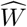 has zero mean. Note that 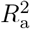 can also be expressed in terms of the Frobenius norm 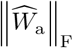 via

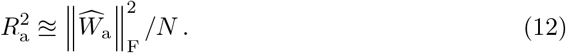

Note that, before proceeding to further investigations, we numerically tested this approximation for *N* = 500 and heterogeneous random sets of *a*_*i*_ drawn from a uniform [0, 1]-distribution and found a very close match to the actual spectral radii (1-2% relative error).

Considering the *R*_a,*i*_ as per site estimates for the spectral radius, one can use the generalized circular law (11) to regulate *R*_a_ on the basis of local adaption rules, one for every *a*_*i*_.

In the particular case of flow control, this was done using a comparison between the variance of neural activity that is present in the network and the resulting recurrent contribution to the membrane potential. A more detailed explanation is given in Sect. *Spectral radius, singular values and global Lyapunov exponents* and S1 Appendix. In short, we propose that,

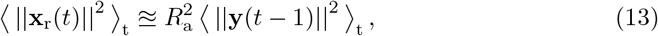

where *x*_r,*i*_ is the recurrent contribution to the membrane potential *x*_*i*_. This stationarity condition led to the adaptation rule given in Eq. (3).

Instead of directly imposing Eq. (13) via an appropriate adaptation mechanism, we also considered the possibility of transferring this condition into a set point for the variance of neural activities as a function the external driving. To do so, we used a mean-field approach to describe the effect of the recurrent input onto the resulting neural activity variance. A detailed description is given in S2 Appendix. This led to the update rule given by Eq. (5)–(9).

## Results

### Flow control

Flow control, see Eq. (3), robustly led to the desired spectral radius *R*_t_ when correlations were weak. It is of particular interest that an internal network parameter, the spectral radius, can be regulated in the continuing present of external inputs **I**(*t*). For correlated input streams, as listed in Sect. *Input protocols*, substantial inter-neuron correlation may be induced. In this case a stationary state was still attained, but with a spectral radius deviating to a certain extent from the parameter *R*_t_ entering (3).

In Fig. 1, we present a simulation for a network with *N* = 500 sites, a connection probability *p*_r_ = 0.1 and an adaption rate *ϵ*_a_ = 10^− 3^. The standard deviation of the external driving is *σ*_ext_ = 0.5. Numerically, we found that the time needed to converge to the stationary states depends substantially on *R*_t_, slowing down when the spectral radius becomes small. It is then advantageous, as we have done, to scale the adaption rate *ϵ*_a_ inversely with the trailing average 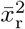 of 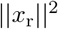, viz as 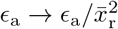.

**Fig 1.**
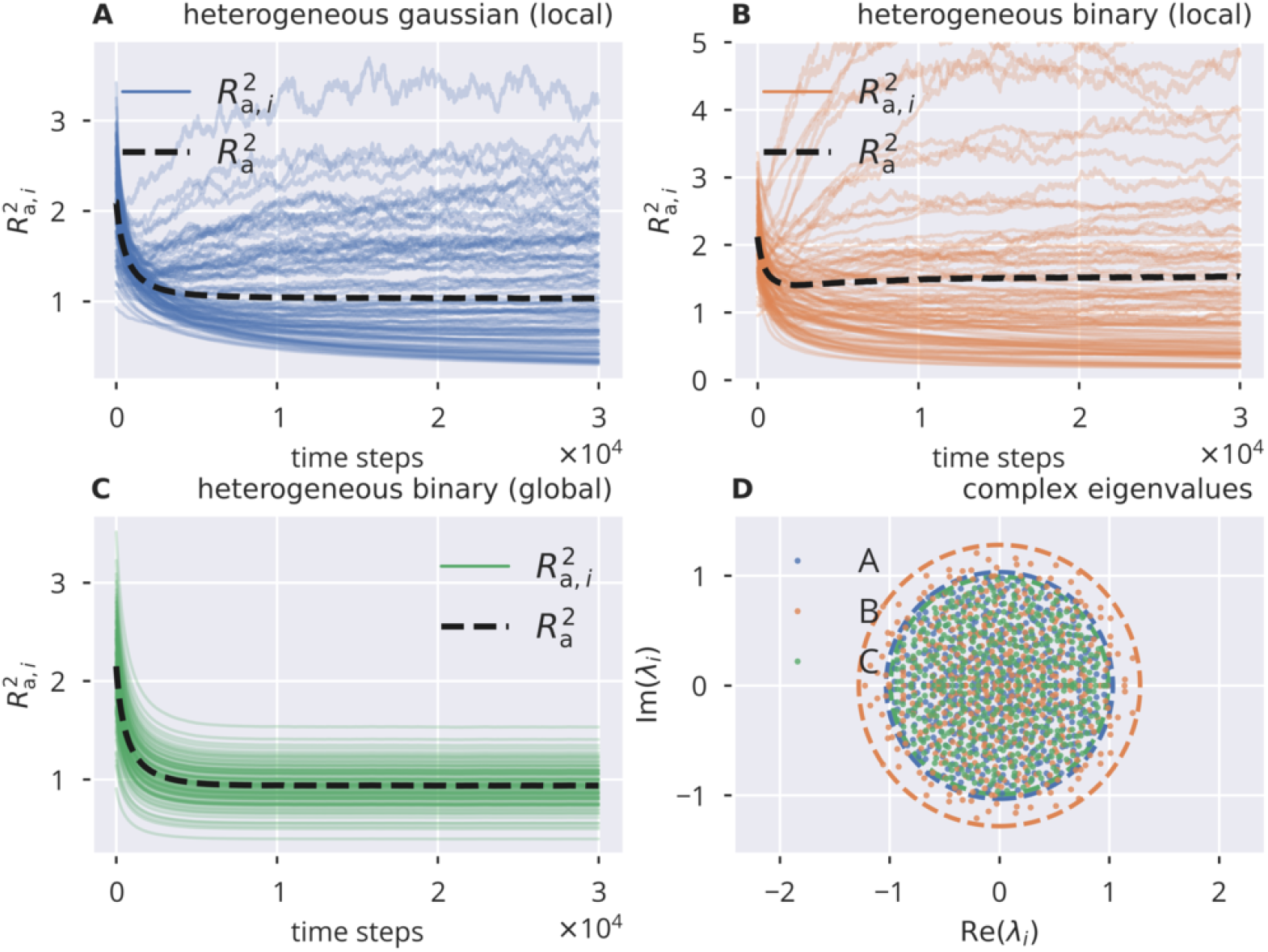
Online spectral radius regulation using flow control. The spectral radius *R*_a_ and the respective local estimates *R*_a,*i*_ as defined by (11). For the input protocols see Sect. *Input protocols*, for the local and global adaption rules (3) and (4). **A**: Dynamics of 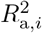 and 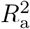, in the presence of heterogeneous independent Gaussian inputs. Local adaptation. **B**: Heterogeneous identical binary input, local adaptation. **C**: Heterogeneous identical binary input, global adaptation. **D**: Distribution of eigenvalues of the corresponding affective synaptic matrices 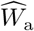. Circles denote the respective spectral radii. Local adaption converges to a spectral radius that is too large (the target is unity), when the driving input is strongly correlated.

For all input protocols and adaption rules, the spectral radius was found to converge to a finite value. The desired value, in this case *R*_t_ = 1, was however not attained when a strongly correlated input interferes with local adaption, see Eq. (3). The local estimates *R*_a,*i*_ for the spectral radius show a substantial heterogeneity, which was reduced when adaption is global, using (16).

### Variance control

In comparison, variance control, shown in Fig. 2, resulted in a spectral radius that was consistently off by a factor of approx. 20%. Note that we had to introduce a lower bound of 0 for *a*_*i*_, since some gain factors would otherwise have become negative. Overall, in comparison to flow control, variance control did not exhibit the same level of precision in tuning the system towards a desired spectral spectral radius.

**Fig 2.**
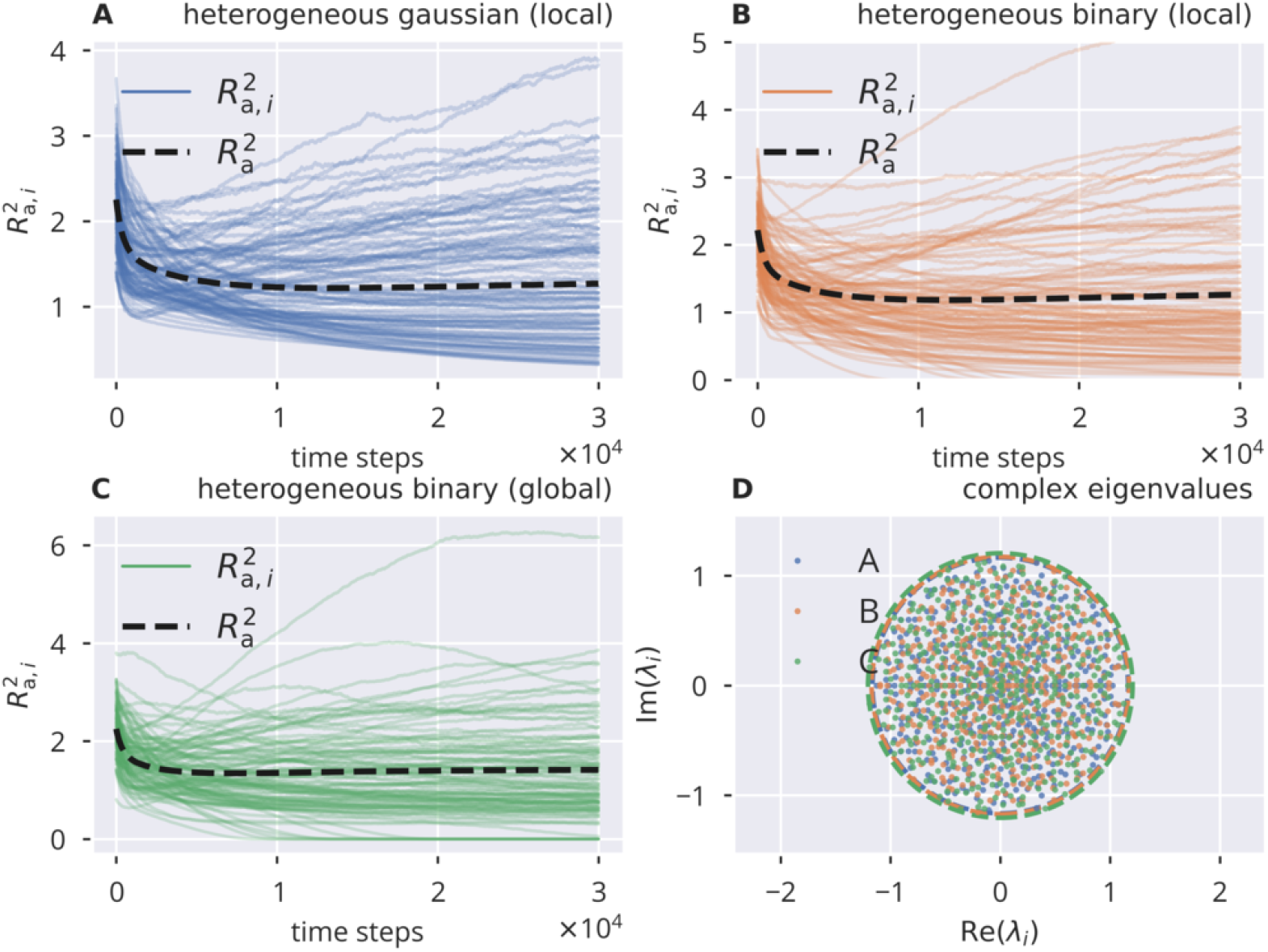
Online spectral radius regulation using variance control. The spectral radius *R*_a_ and the respective local estimates *R*_a,*i*_ as defined by (11). For the input protocols see Sect. *Input protocols*, for the local and global adaption rules (5) and (10). **A**: Dynamics of 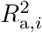 and 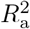, in the presence of heterogeneous independent Gaussian inputs. Local adaptation. **B**: Heterogeneous identical binary input, local adaptation. **C**: Heterogeneous identical binary input, global adaptation. **D**: Distribution of eigenvalues of the corresponding affective synaptic matrices 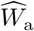. Circles denote the respective spectral radii. Local adaption converges to a spectral radius that is too large (the target is *R*_t_ = 1) under all input protocols.

### Spectral radius, singular values and global Lyapunov exponents

Apart from the spectral radius *R*_a_ of the matrix 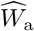, one may consider the relation between the adaptation dynamics and the respective singular values *σ*_*i*_ of 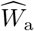. We recall that the spectrum of 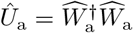 is given by the squared singular values, 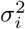, and that the relation 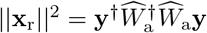 holds. Now, assume that the time-averaged projection of neural activity **y** = **y**(*t*) onto all eigenvectors of *U* (*t*) is approximately the same, that is, there is no preferred direction of neural activity in phase space. From this idealized case, it follows that the time average of the recurrent contribution to the membrane potential can be expressed with

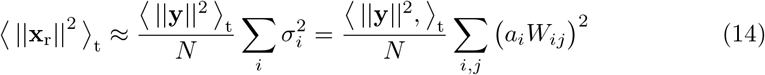

as the rescaled average of the 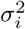. For the second equation, we used the fact that the 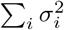 equals the sum of all matrix elements squared [36, 37]. With (11), one finds that (14) is equivalent to 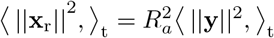 and hence to the flow condition (13). This result can be generalized, as done in S1 Appendix, to the case that the neural activities have inhomogeneous variances, while still being uncorrelated with zero mean. We then show that the stationarity condition leads to a spectral radius of (approximately) unity.

It is worthwhile to note that the singular values of 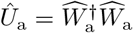 do exceed unity when *R*_a_ = 1. More precisely, for a random matrix with i.i.d. entries, one finds in the limit of large *N* that the largest singular value is given by *σ*_max_ = 2*R*_a_, in accordance with the Marchenko-Pastur law for large random matrices [38]. Consequently, directions in phase space exist, in which the norm of the phase space vector is elongated by factors greater than one. Still, this does not contradict the fact that a unit spectral radius coincides with a transition to chaos for the non-driven case. The reason is that the global Lyapunov exponents are given by

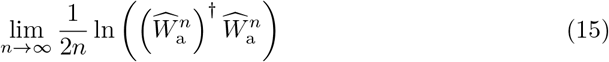

which eventually converge to ln ‖*λ*‖, see S1 Fig and [39], where *λ*_*i*_ is the *i*th eigenvalue of 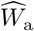. The largest singular value of the *n*th power of a random matrix with a spectral radius *R*_a_ scales like 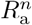 in the limit of large powers *n*. The global Lyapunov exponent goes to zero as a consequence when *R*_a_ → 1.

### Spectral radius adaption dynamics

For an understanding of the spectral radius adaption dynamics of flow control, it is of interest to examine the effect of using the global adaption constraint

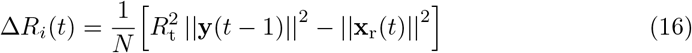

in (3). The spectral radius condition (13) is then enforced directly, with the consequence that (16) is stable and precise even in the presence of correlated neural activities (see Fig. 1C). This rule, while not biologically plausible, provides an opportunity to examine the dynamical flow, besides the resulting state. There are two dynamical variables, *a* ≡ *a*_*i*_ = *a*_*j*_, which, for the sake of simplicity, are assumed to be homogeneous, and the activity variance 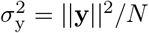. The evolution of 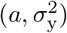 resulting from the global rule (4) is shown in Fig. 3.

**Fig 3.**
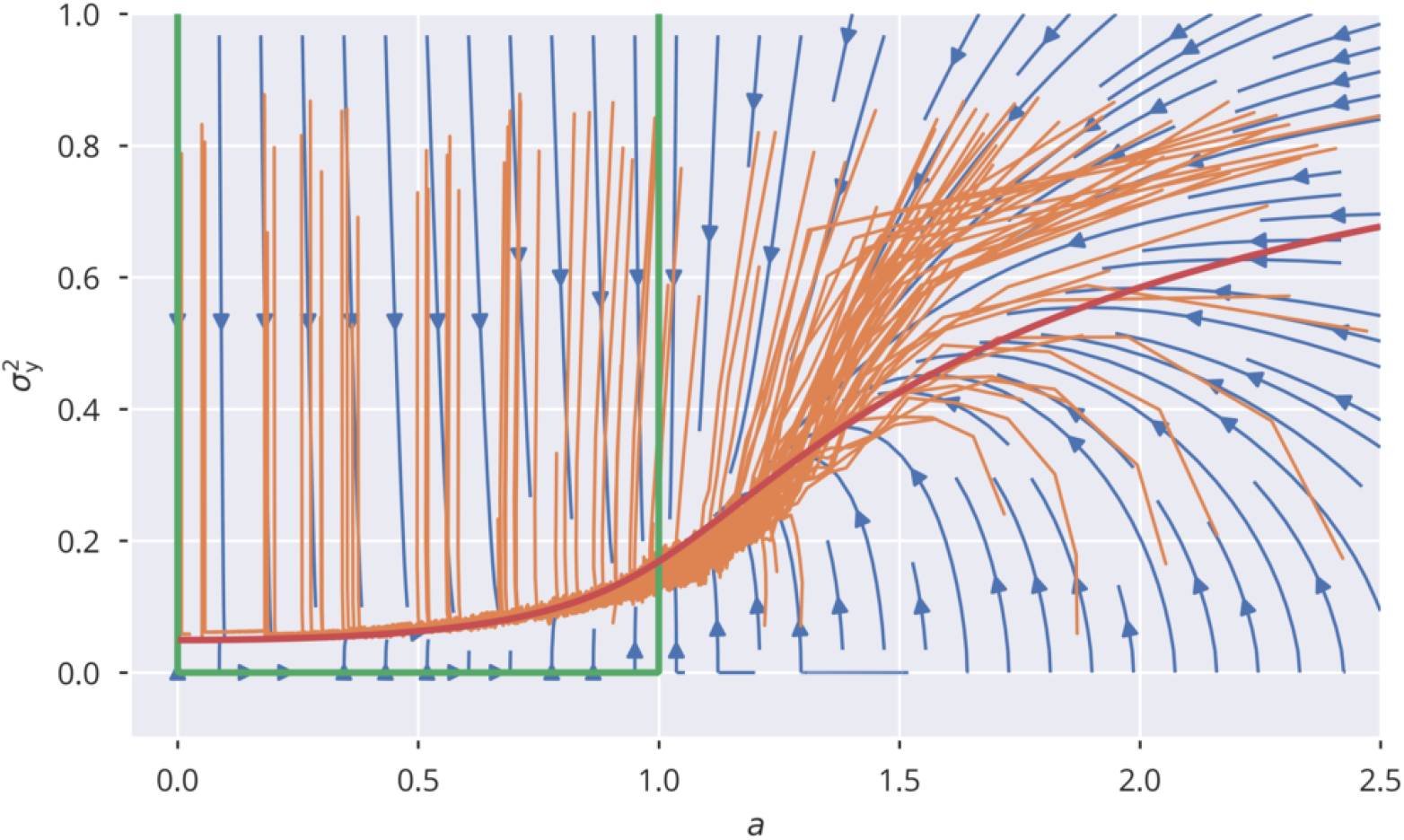
Spectral radius adaption dynamics. The dynamics of the synaptic rescaling factor *a* and the squared activity 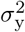 (orange), as given by (4), for *R*_t_ = 1. Also shown is the analytic approximation to the flow (blue), see (17) and (18), and the respective nullclines Δ*a* = 0 (green) and 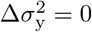 (red). For the input, the heterogeneous binary protocol is used, with *σ*_ext_ = 0.25 and *ϵ*_a_ = 0.1.

For the flow, Δ*a* = *a*(*t* + 1) − *a*(*t*) and 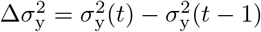, the approximation

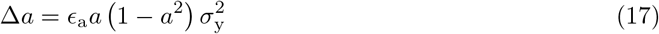

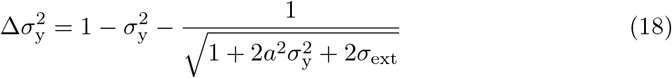

is obtained when using *R*_t_ = 1 in (16), and *σ*_w_ = 1 for the normalized variance of the synaptic weights, as defined by (26). We used the mean-field approximation for neural variances that is derived in in S2 Appendix. The analytic flow compares well with numerics, as shown in Fig. 3. For a subcritical rescaling factor *a* and *σ*_ext_ = 0, the system flows towards a line of fixpoints defined by a vanishing 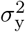 and a finite *a* ∈ [0, 1]. When starting with *a* > 0, the fixpoint is instead 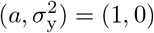. The situation changes qualitatively for finite external inputs, viz when *σ*_ext_ > 0. The nullcline 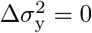 is now continuous and the system flows, as shown in Fig. 3 to *a* = 1, with the value of 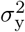 being determined by the intersection of the two nullclines.

This analysis shows that external input is necessary for a robust flow towards the desired spectral weight, the reason being that the dynamics dies out before the spectral weight can be adapted when the isolated systems starts in the subcritical regime.

### XOR-memory recall

To this point, we only presented results regarding the effectiveness of the introduced adaption rules. However, we did not account for their effects onto a given learning task. Therefore, we tested the performance of locally adapted networks under the delayed XOR task, which evaluates the memory capacity of the echo state network in combination with a non-linear operation. For the task, the XOR operation is to be taken with respect to a delayed pair of two consecutive binary inputs signals, *u*(*t* − *τ*) and *u*(*t* − *τ* − 1), where *τ* is a fixed time delay. The readout layer is given by a single unit, which has the task to reproduce

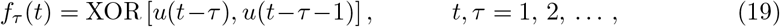

where XOR[*u*, *u*′] is 0/1 if *u* and *u*′ are identical/not identical.

The readout vector **w**_out_ is trained with respect to the mean squared output error,

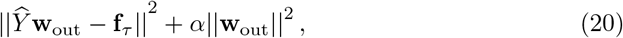

using ridge regression on a sample batch of *T*_batch_ = 10*N* time steps, here for *N* = 500, and a regularization factor of *α* = 0.01. The batch matrix 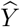, of size *T*_batch_ × (*N* + 1), holds the neural activities as well as one node with constant activity serving as a potential bias. The readout (column) vector **w**_out_ is similarly of size (*N* + 1). The *T*_batch_ entries of **f**_*τ*_ are the *f*_*τ*_ (*t*), viz the target values of the XOR problem. Minimizing (20) leads to

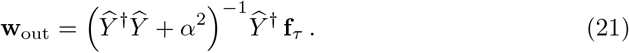

The learning procedure is repeated independently for each time delay *τ*. We quantified the performance by the total memory capacity, MC_XOR_, as

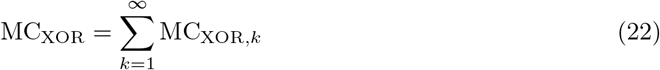

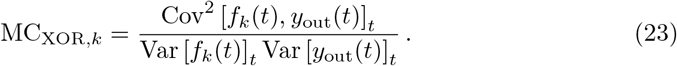

The activity 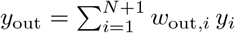 of the readout unit is compared in (23) with the XOR prediction task, with the additional neuron, *y*_*N*+1_ = 1, corresponding to the bias of the readout unit. The variance and covariance are calculated with respect to the batch size *T*_batch_.

The results for flow control presented in Fig. 4 correspond to two input protocols, heterogeneous Gaussian and binary inputs. Shown are sweeps over a range of *σ*_ext_ and *R*_t_. The update rule (3) was applied to the network for each pair of parameters until the *a*_*i*_ values converged to a stable configuration. We then measured the task performance as described above. Note that in the case of Gaussian input, this protocol was only used during the adaptation phases. Due to the nature of the XOR task, we chose to use binary inputs with the corresponding variances during the performance testing. See S2 Fig in the appendix for a performance sweep using the homogeneous binary and Gaussian input protocol. Optimal performance is generally attained around the *R*_a_ ≈ 1 line. A spectral radius *R*_a_ slightly smaller than unity was optimal when using Gaussian input, but not for binary input signals. In this case the measured spectral radius *R*_a_ deviates linearly from the target *R*_t_, with increasing strength of the input, as parameterized by the standard deviation *σ*_ext_. Optimal performance is, however, essentially independent of the input strength, apart from *σ*_ext_ → 0, with the peak located roughly at *R*_t_ ≈ 0.55.

**Fig 4.**
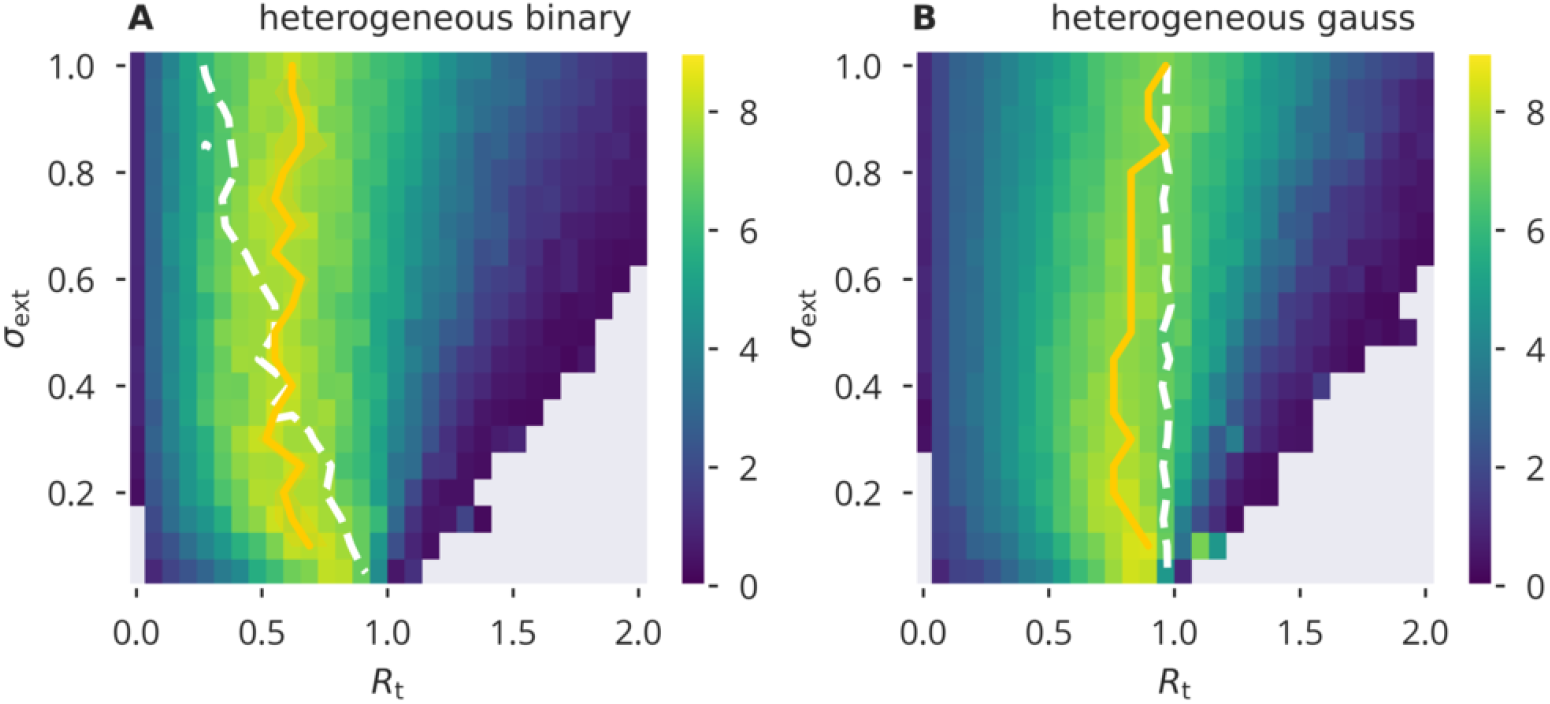
XOR performance for flow control. Color-coded performance sweeps for the XOR-performance (22) after adaptation using flow control. Averaged over five trials. The input has variance 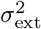 and the target for the spectral radius *R*_t_. A/B panels are for heterogeneous binary/Gaussian input protocols. Optimal performance (yellow solid line) is in general close to criticality, *R*_a_ = 1, as measured (white dashed lines).

Comparing these results to variance control, as shown in Fig. 5, we found that variance control resulted in an overall lower performance. To our surprise, for external input with a large variance, Gaussian input caused stronger deviations from the desired spectral radius as compared to binary input. Therefore, in a sense, it appeared to behave opposite to what we found for flow control. However, similar to flow control, the value of *R*_t_ giving optimal performance under a given *σ*_ext_ remained relatively stable over the range of external input strength measured. On the other hand, using homogeneous input, see S3 Fig, did cause substantial deviations from the target spectral radius when using binary input.

**Fig 5.**
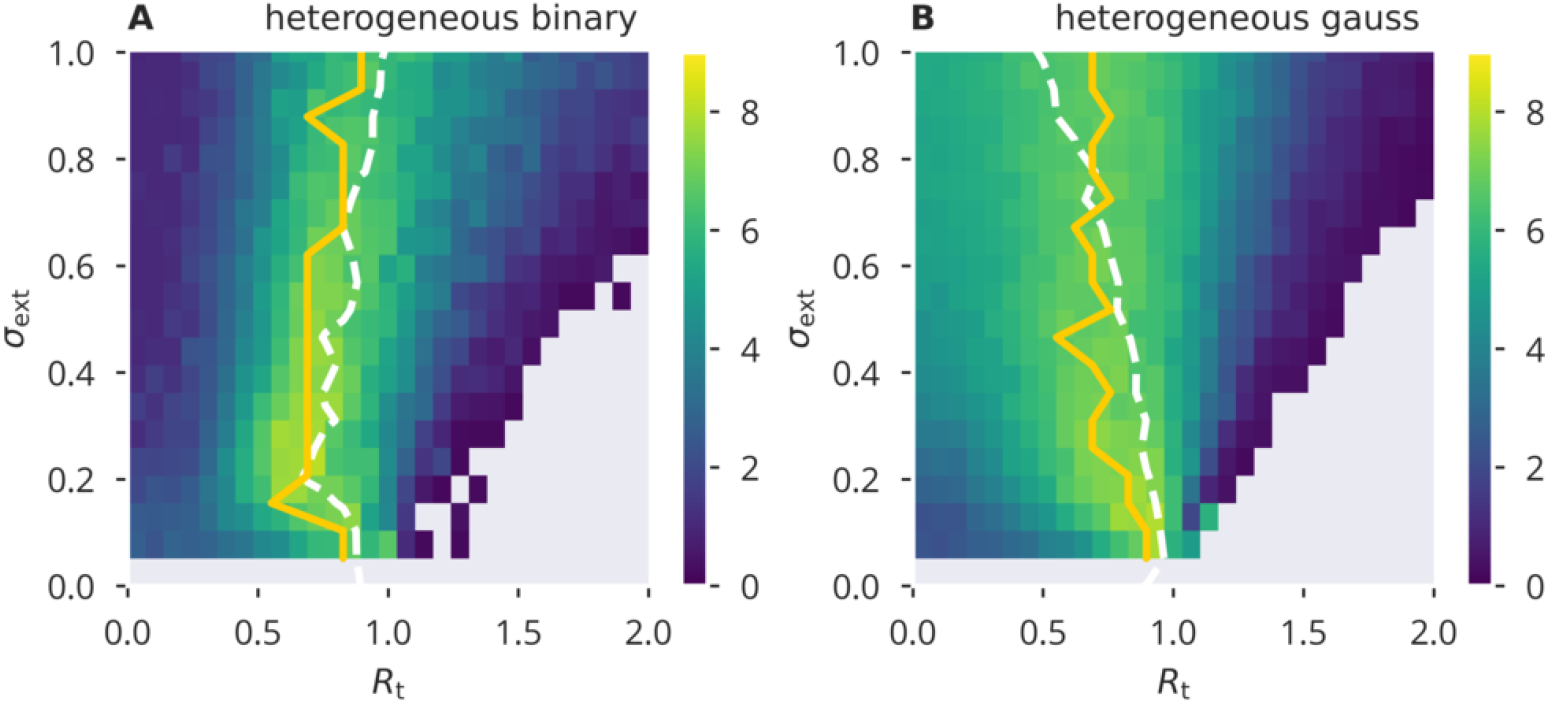
XOR performance for variance control. Color-coded performance sweeps for the XOR-performance (22) after adaptation using variance control. Averaged over five trials. The input has variance 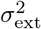 and the target for the spectral radius *R*_t_. A/B panels are for heterogeneous binary/Gaussian input protocols. Optimal performance (yellow solid line) is in general close to criticality, *R*_a_ = 1, as measured (white dashed lines).

The two input protocols used for the data shown in Fig. 4 and 5 relate to two different (biological) scenarios: First, an idealized case where a local recurrent neural ensemble only receives input from one particular source that has very specific statistics, such as, in our model case, a binary input sequence. In this scenario, it is plausible to assume that the input statistics under which long term adaptation is taking place coincide with the type of input patterns that are considered for the evaluation of the actual task performance. A second, presumably more plausible situation is the case where different input streams are integrated in the neural ensemble. Each neuron receives a weighted superposition of those input signals. Unless those inputs were strongly correlated, this would mean that random Gaussian input—as used in Fig. 4 B and 5 B—is a reasonable approximation to this scenario.

### Input induced correlations

The theory for the echo-state layer, Eq. (49), which was used in Eq. (18) and (6), assumes that the variance 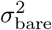 of the bare recurrent contribution to the membrane potential, *x*_bare_ = ∑_*j*_ *W*_*ij*_*y*_*j*_ is given by 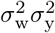 In Fig. 6, a comparison is presented for the four input protocols introduced in Sect. *Input protocols*. For the Gaussian protocols, for which neurons receive statistically uncorrelated external signals, one observes that 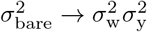 in the thermodynamic limit *N* → ∞ via a power law, which is to be expected when the presynaptic neural activities are decorrelated. On the other side, binary 0/1 inputs act synchronous on all sites, either with site-dependent or site-independent strengths (heterogeneous/homogeneous). Corresponding activity correlations are induced and a finite and only weakly size-dependent difference between 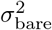 and 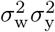 shows up. Substantial corrections to the analytic theory are to be expected in this case. To this extend we measured the cross-correlation *C*(*y*_*i*_, *y*_*j*_), defined as

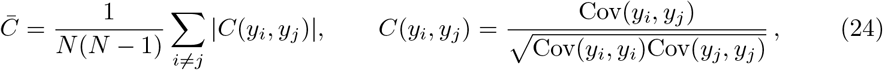

with the covariance given by Cov(*y*_*i*_, *y*_*j*_) = 〈(*y*_*i*_ − 〈*y*_*i*_〉_*t*_)(*y*_*j*_ − 〈*y*_*j*_〉_*t*_)〉_*t*_. For a system of *N* = 500 neurons the results for the averaged absolute correlation 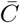 are presented in Fig. 7 (see S4 Fig in the supplementary material for homogeneous input protocols). Autonomous echo-state layers are in chaotic states when supporting a finite activity level, which implies that correlations vanish in the thermodynamic limit *N* → ∞. The case *σ*_ext_ = 0, as included in Fig. 7, serves consequently as a yardstick for the magnitude of correlations that are due to the finite number of neurons.

**Fig 6.**
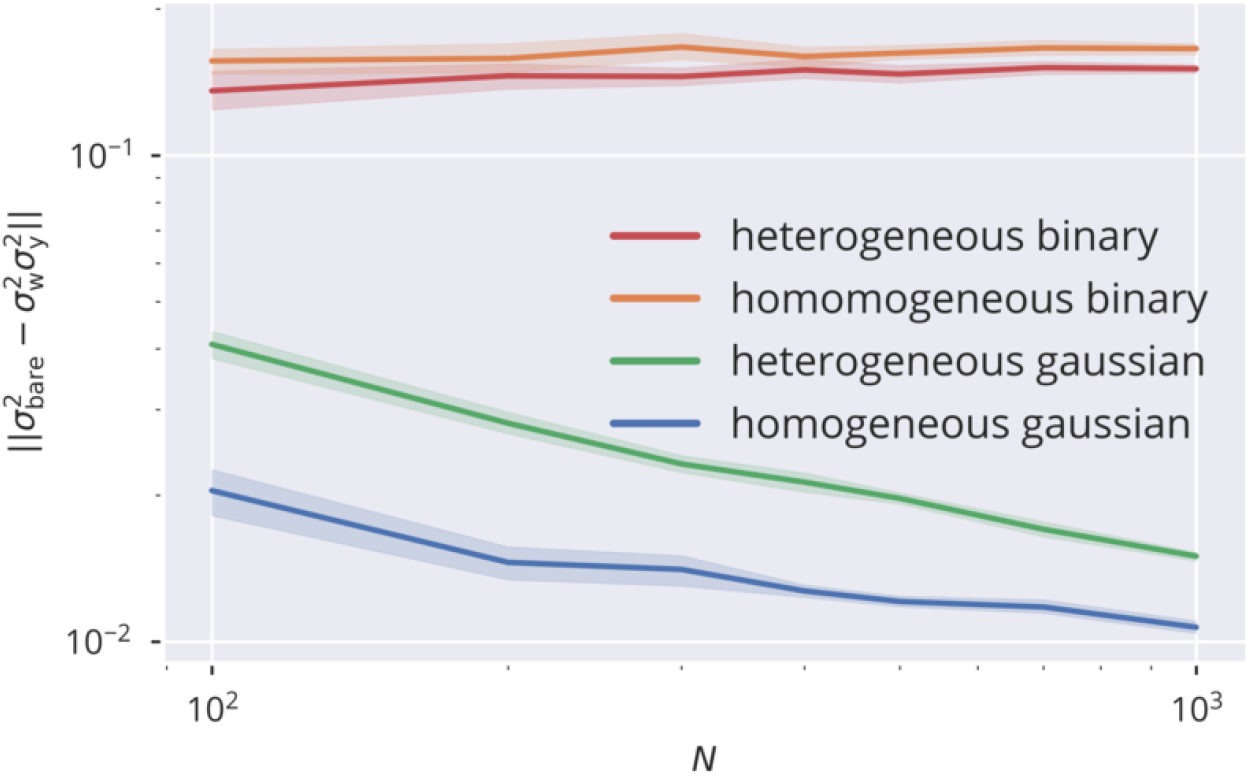
Size dependence of correlation. Comparison between the variance 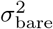 of the bare recurrent input *x*_bare_ = ∑_*j*_ *W*_*ij*_*y*_*j*_ with 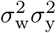 Equality is given when the presynaptic activities are statistically independent. This can be observed in the limit of large network sizes *N* for uncorrelated input data streams (homogeneous and heterogeneous Gaussian input protocols), but not for correlated inputs (homogeneous and heterogeneous binary input protocols). Compare Sect. *Input protocols* for the input protocols. Parameters are *σ*_ext_ = 0.5, *R*_a_ = 1 and *μ*_*t*_ = 0.05.

**Fig 7.**
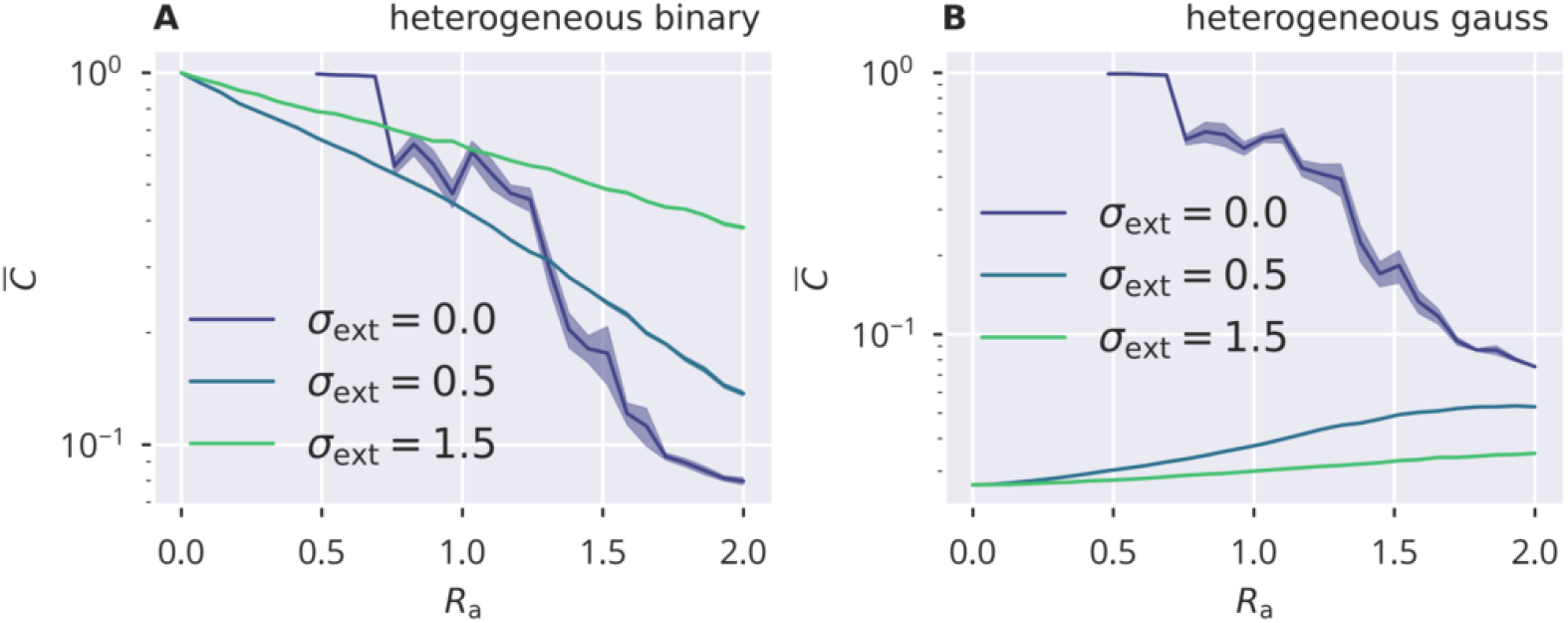
Input induced activity correlations. For heterogeneous binary and Gaussian inputs (A/B), the dependency of mean activity cross correlations 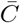, see Eq. (24). 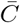 is shown as a function of the target spectral radius *R*_a_. Results are obtained for *N* = 500 sites by averaging over five trials, with shadows indicating the accuracy. Correlations are due to finite-size effect for the autonomous case *σ*_ext_ = 0.

**Fig 8.**
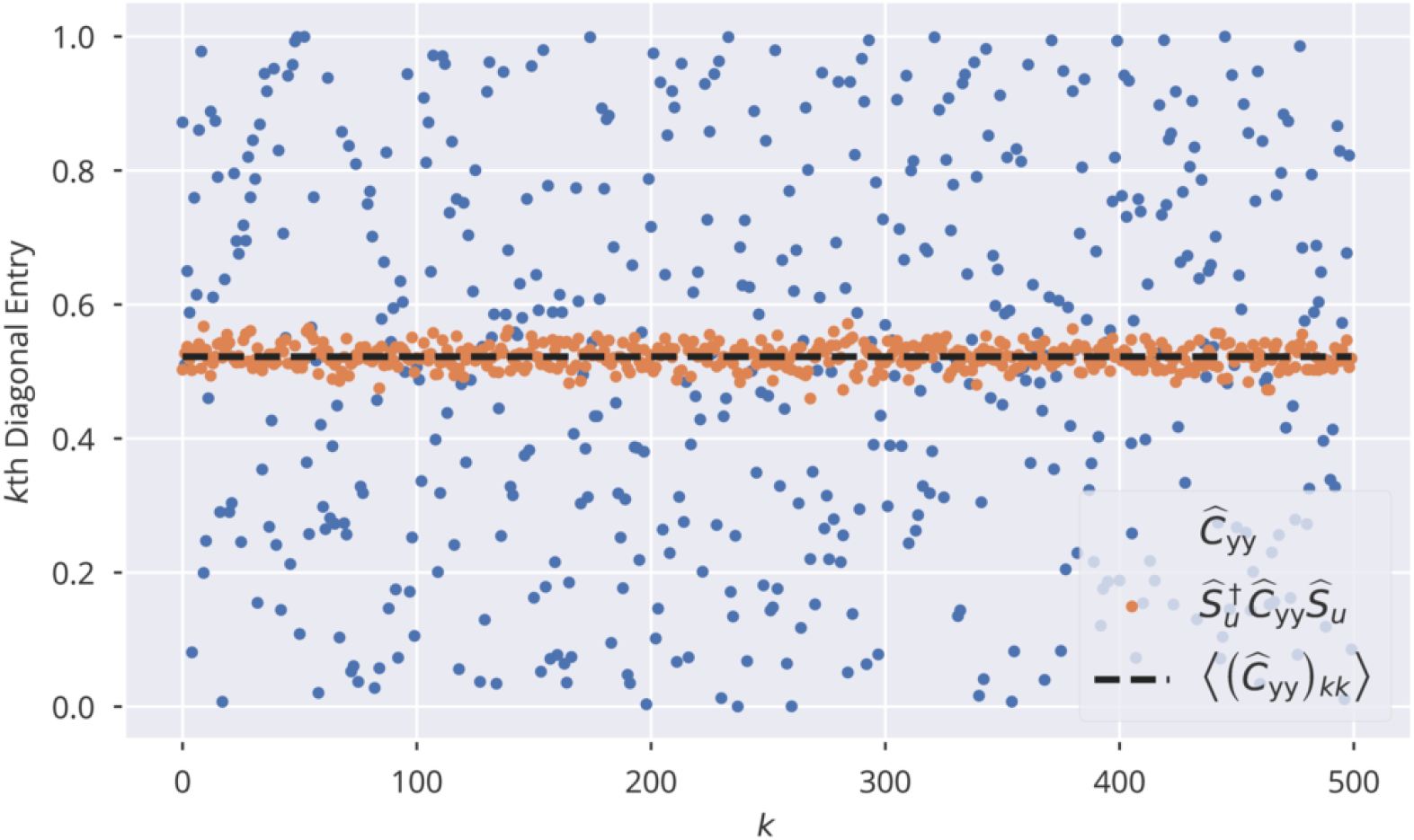
Diagonal Elements of a randomly generated covariance matrix and its representation in the 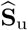 basis. 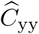 is a diagonal matrix with diagonal entries randomly drawn from [0, 1], 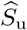 is the orthonormal eigenbasis of 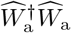, where 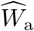 is a random Gaussian matrix. The black dashed line denotes the average over the diagonal entries of 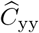.

Input correlations are substantially above the autonomous case for correlated binary inputs, with the magnitude of 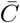 decreasing when the relative contribution of the recurrent activity increases. This is the case for increasing *R*_t_. The effect is opposite for the Gaussian protocol, for which the input does not induce correlations, but contributes to decorrelating neural activity. The mean absolute correlation 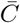 is in this case suppressed when the internal activity becomes small in the limit *R*_t_ → 0. Finite-size effects are relevant only when the internal activity is of the order of the input-induced activity, viz for increasing *R*_t_.

## Discussion

The mechanisms for tuning the spectral radius via a local homeostatic adaption rule introduced in the present study require neurons to have the ability to distinguish and measure locally both external and recurrent input contributions. For flow control, neurons need to be able to compare the recurrent membrane potential with their own activity, as assumed in Sect. *Autonomous spectral radius regulation*. On the other hand, variance control directly measures the variance of the external input and derives the activity target variance accordingly. The limiting factor to a successful spectral radius control is the amount of cross-correlation induced by external driving statistics. As such, the functionality and validity of the proposed mechanisms depended on the ratio between external input, i.e. feed-forward or feedback connections, with respect to recurrent, or lateral connections. In general, it is not straightforward to directly connect experimental evidence regarding the ratio between recurrent and feed-forward contributions to the effects observed in the model. It is, however, worthwhile to note that the fraction of synapses associated with interlaminar loops and intralaminar lateral connections are estimated to make up roughly 50% [40]. Relating this to our model, it implies that the significant interneural correlations that we observed when external input strengths were of the same order of magnitude as the recurrent inputs, can not generally be considered an argifact of biologically implausible parameter choices. In fact, synchronization is a widely observed phenomenon in the brain and is even considered important for information processing [41, 42].

Overall, we found flow control to be generally more robust than variance control in the sense that, while still being affected by the amount of correlations within the neural reservoir, the task performance was less so prone to changes in the external input strength. Comparatively stable network performance could be observed, in spite of significant deviations from the desired spectral radius (see Fig. 4). A possible explanation may be that flow control uses a distribution of samples from only a restricted part of phase space, that is, from the phase space regions that are actually visited or “used” for a given input. Therefore, while a spectral radius of unity ensures–statistically speaking– the desired scaling properties in all phase-space directions, it seem to be enough to control the correct scaling for the subspace of activities that is actually used for a given set of input patters. Variance control, on the other hand, relies more strictly on the assumptions regarding the statistical independence of neural activities. In consequence, the desired results could only be achieved under a rather narrow set of input statistics (independent Gaussian input with small variance). In addition, the approximate expression derived for the nonlinear transformation appearing in the mean field approximation adds another layer of potential source of systematic error to the control mechanism. This aspect also speaks in favor of flow control, since its rules are mathematically more simple. In contrast to variance control, the stationarity condition stated in Eq. (13) is independent of the actual nonlinear activation function used and could easily be adopted in a modified neuron model. It should be noted, however, that the actual target *R*_t_ giving optimal performance might then also be affected.

Interestingly, flow control distinguishes itself from a conventional local activity-target perspective of synaptic homeostasis: There is no predefined set point in Eq. (3). This allows heterogeneities of variances of neural activity to develop across the network, while retaining the average neural activity at a fixed predefined level.

## Conclusion

Apart from being relevant from a theoretical perspetive, we propose that the separability of recurrent and external contributions to the membrane potential is an aspect that is potentially relevant for the understanding of local homeostasis in biological networks. While homeostasis in neural compartments has been a subject of experimental research [43, 44], to our knowledge, it has not yet been further investigated on a theoretical basis. Given that the neural network model used in this study lacks some features characterizing biological neural networks (e.g. strict positivity of the neural firing rate, Dale’s law), future research should therefore investigate whether the here presented framework of local flow control can be implemented within more realistic biological neural network models. A particular concern regarding our findings is that biological neurons are spiking. The concept of an underlying instantaneous firing rate is, strictly speaking, a theoretical construct, let alone the definition of higher moments, such as the “variance of neural activity”. However, it is important to note that real-world biological control mechanims, e.g. of the activity, rely on physical quantities that serve as measurable correlates. A well-known example is the intracellular calcium concentration, which is essentially a linearly filtered version of the neural spike train [32]. On a theoretical level, Cannon and Miller showed that dual homeostasis can successfully control the mean and variance of this type of spike-averaging physical quantities [26]. An extension of the flow control to filtered spike trains of spiking neurons could be an interesting subject of further investigations. However, using spiking neuron models would have shifted the focus of our research towards the theory of liquid state machines [45, 46], exceeding the scope of this publication. We therefore leave the extension to more realistic network/neuron models to future work.

## Materials and methods

### Model

We implemented an echo state network with *N* neurons, receiving *D*_in_ inputs. The neural activity is *y*_*i*_ ∈ [−1, 1], *x*_*i*_ the membrane potential, *u*_*i*_ the input activities, *W*_*ij*_ the internal synaptic weights and *I*_*i*_ the external input received. The output layer will be specified later. The dynamics

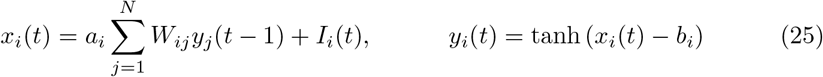

is discrete in time, where the input *I*_*i*_ is treated instantaneously. A tanh-sigmoidal has been used as a nonlinear activation function.

The synaptic renormalization factor *a*_*i*_ in (25) can be thought of as a synaptic scaling parameter that neurons use to regulate the overall strength of the recurrent inputs. The strength of the inputs *I*_*i*_ is unaffected, which is biologically plausible if external and recurrent signals arrive at separate branches of the dendritic tree [44].

The *W*_*ij*_ are the bare synaptic weights, with *a*_*i*_*W*_*ij*_ being the components of the effective weight matrix 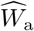. Key to our approach is that the propagation of activity is determined by 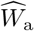, which implies that the spectral radius of the effective, and not of the bare weight matrix needs to be regulated.

The bare synaptic matrix *W*_*ij*_ is sparse, with a connection probability *p*_r_ = 0.1. The non-zero elements are drawn from a Gaussian with standard deviation

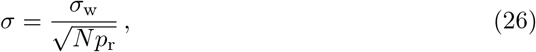

and vanishing mean *μ*. Here 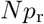 corresponds to the mean number of afferent internal synapses, with the scaling 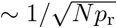 enforcing size-consistent synaptic-weight variances.

As introduced in the introduction, we applied the following adaptation mechanisms:

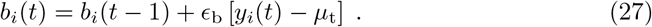

For the gains, using flow control:

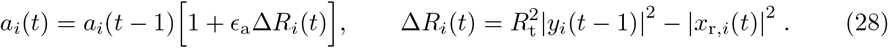

For variance control:

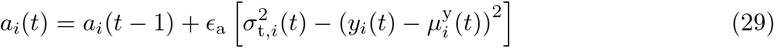

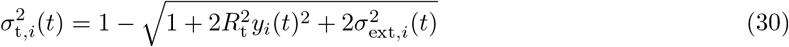

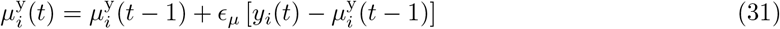

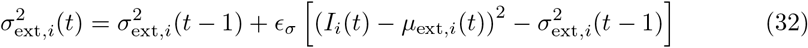

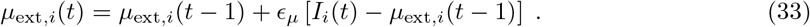

For a summary of model parameters, see Table 1.

**Table 1.**
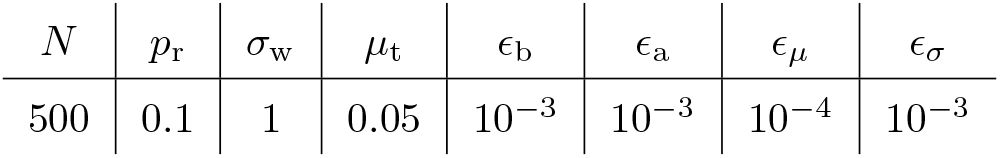
Standard values for model parameters

### Input protocols

Overall, we examined four distinct input protocols.

1. *Homogeneous Gaussian.* Nodes receive inputs *I*_*i*_(*t*) that are drawn individually from a Gaussian with vanishing mean and standard deviation *σ*_ext_.
2. *Heterogeneous Gaussian.* Nodes receive stochastically independent inputs *I*_*i*_(*t*) that are drawn from Gaussian distributions with vanishing mean and node specific standard deviations *σ*_*i*,ext_. The individual *σ*_*i*,ext_ are normal distributed, as drawn from the positive part of a Gaussian with mean zero and variance 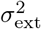.
3. *Homogeneous binary.* Sites receive identical inputs *I*_*i*_(*t*) = *σ*_ext_*u*(*t*), where *u*(*t*) = ±1 is a binary input sequence.
4. *Heterogeneous binary.* We define with

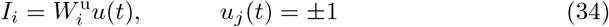

the afferent synaptic weight vector 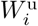, which connects the binary input sequence *u*(*t*) to the network. All 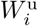 are drawn independently from a Gaussian with mean zero and standard deviation *σ*_ext_.

The Gaussian input variant simulates external noise. We used it in particular to test predictions of the theory developed in Sect. *S2 Appendix*. In order to test the performance of the echo state network with respect to the delayed XOR task, the binary input protocols are employed. A generalization of the here defined protocols to the case of vectorial input signals would be straightforward.

**Fig 9.**
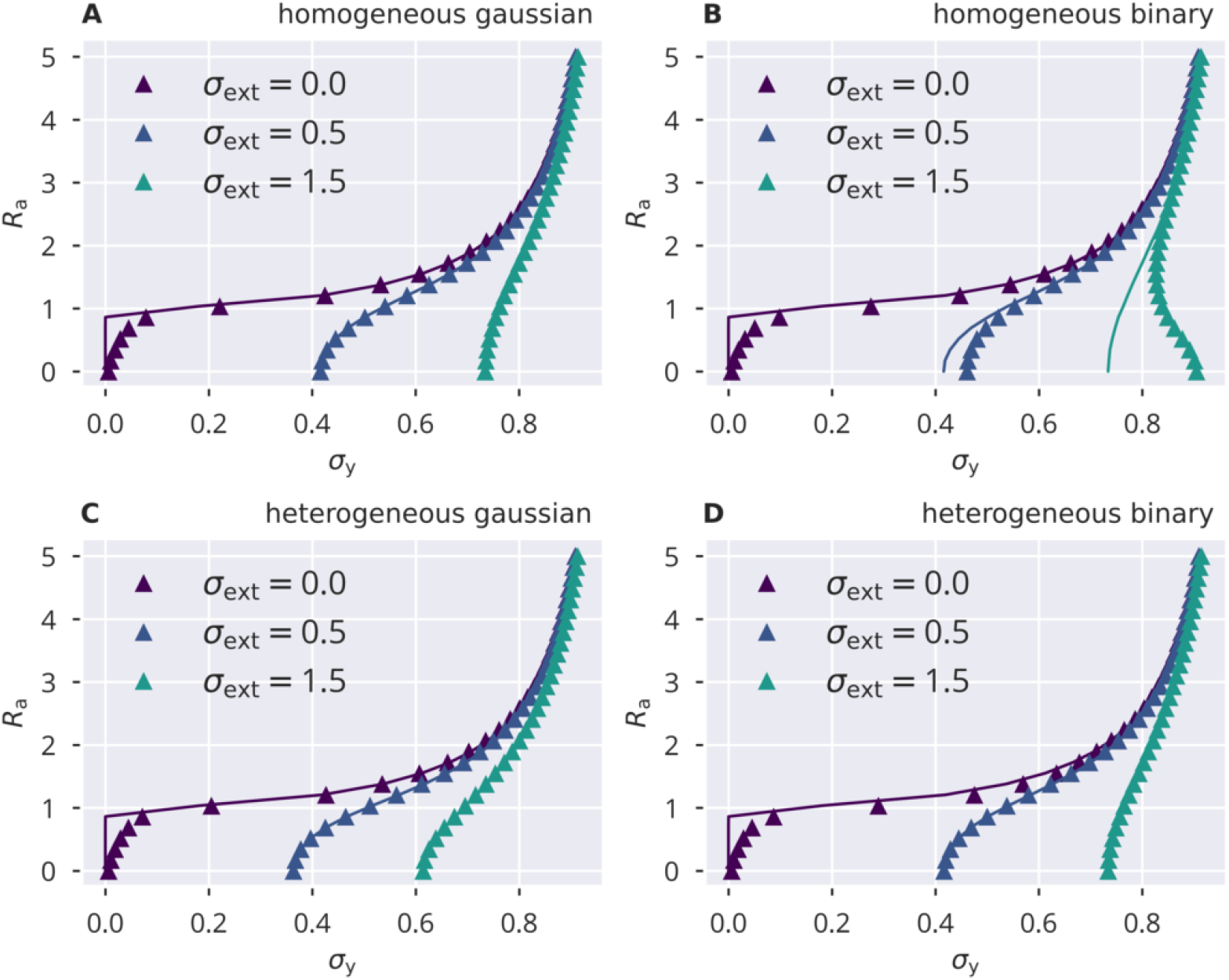
Variance control for the spectral radius. The spectral radius *R*_a_, given by the approximation 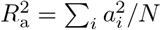, for the four input protocols defined in Sect. *Input protocols*. Lines show the numerical self-consistency solution of (49), symbols the full network simulations. Note the instability for small *σ*_y_ and *σ*_ext_. **A**: Homogeneous independent Gaussian input. **B**: Homogeneous identical binary input. **C**: Heterogeneous independent Gaussian input. **D**: Heterogeneous identical binary input.

## Supporting information

**S1 Fig.**
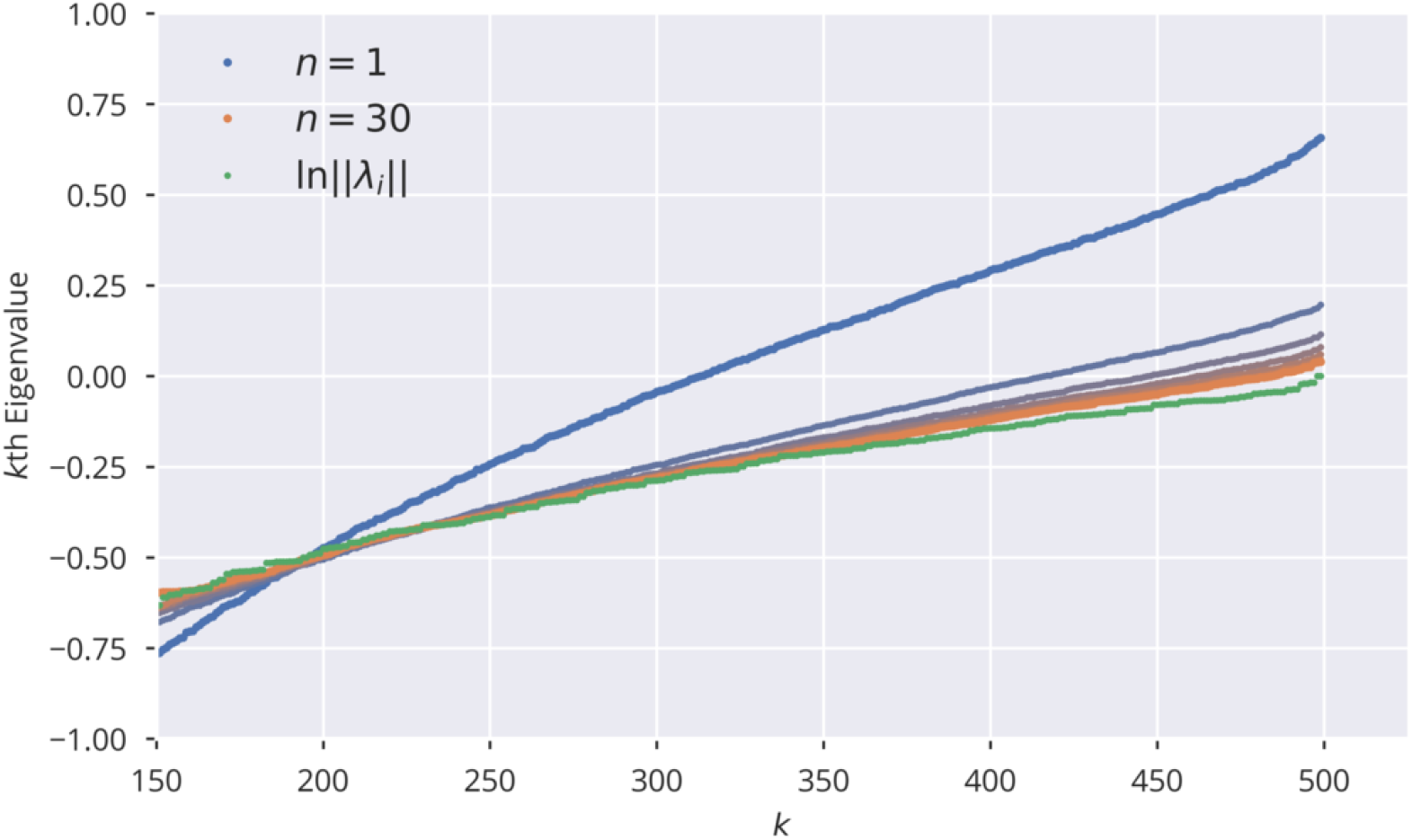
Convergence of Lyapunov Spectrum. Convergence of eigenvalues of 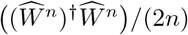 for different *n*, as discussed in Sect.. 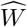 is a random Gaussian matrix which was rescaled to a spectral radius of one. Colors from blue to orange encode the exponent *n* ranging between 1 and 30. Green dots show the theoretical limit of ln‖*λ*_*i*_‖, where *λ*_*i*_ is the *i*th eigenvalue of 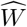.

**S2 Fig.**
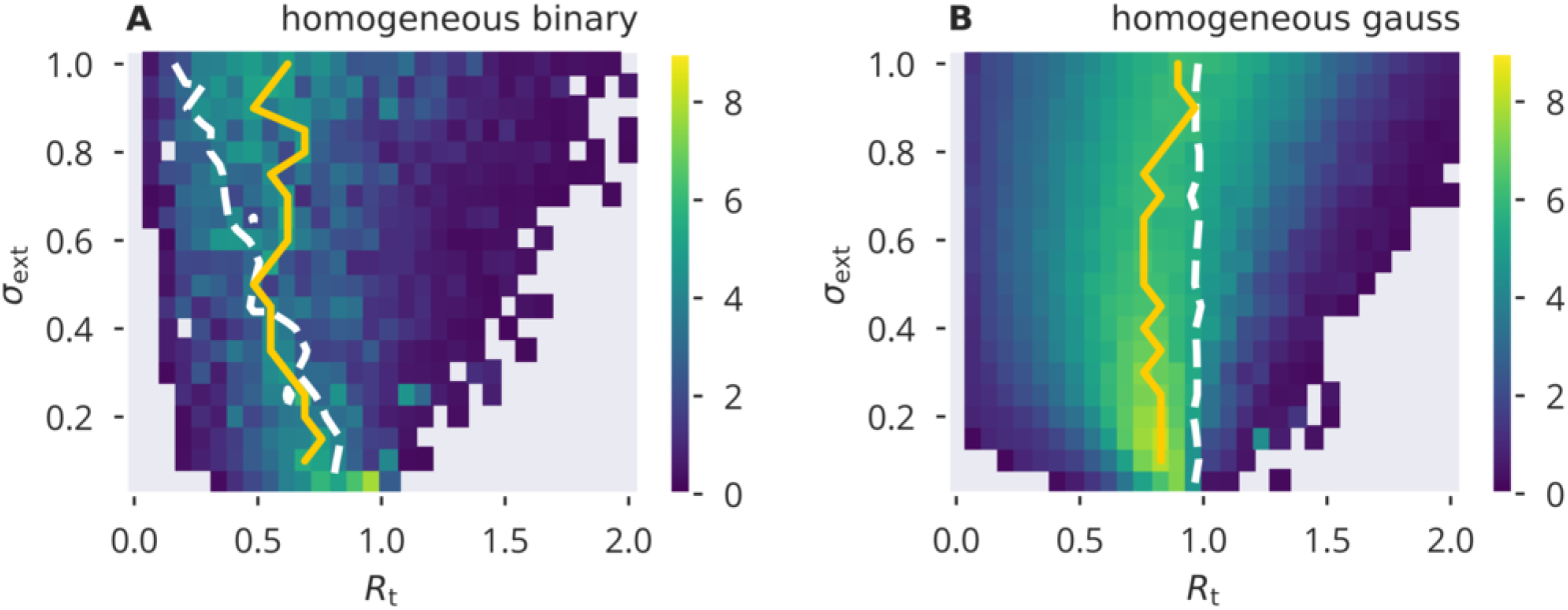
XOR performance for flow control, homogeneous input. Numerical results for the network performance under a time-delayed XOR task, as defined in Sect. *XOR-memory recall*, using homogeneous binary/Gaussian input. Shown are color-coded performance sweeps for the XOR-performance (22), averaged over five trials. The input has variance 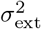 and the target for the spectral radius *R*_t_. A/B panels are for binary/Gaussian input protocols. Optimal performance for a given *σ*_ext_ is given by yellow solid lines, measured value of *R*_a_ = 1 by white dashed lines.

**S3 Fig.**
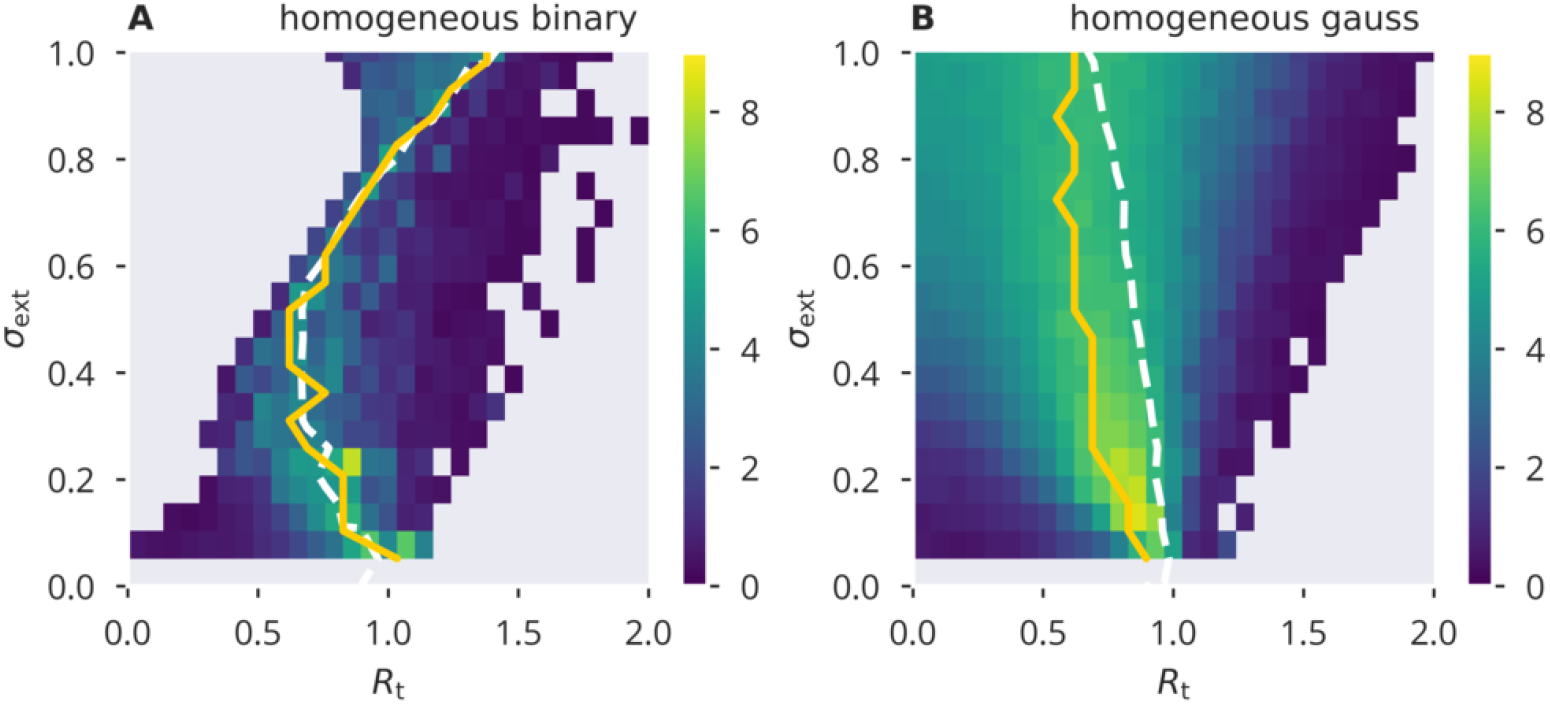
XOR performance for variance control, homogeneous input. Numerical results for the network performance under a time-delayed XOR task, as defined in Sect. *XOR-memory recall*, using homogeneous binary/Gaussian input. Shown are color-coded performance sweeps for the XOR-performance (22), averaged over five trials. The input has variance 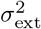 and the target for the spectral radius *R*_t_. A/B panels are for binary/Gaussian input protocols. Optimal performance for a given *σ*_ext_ is given by yellow solid lines, measured value of *R*_a_ = 1 by white dashed lines.

**S4 Fig.**
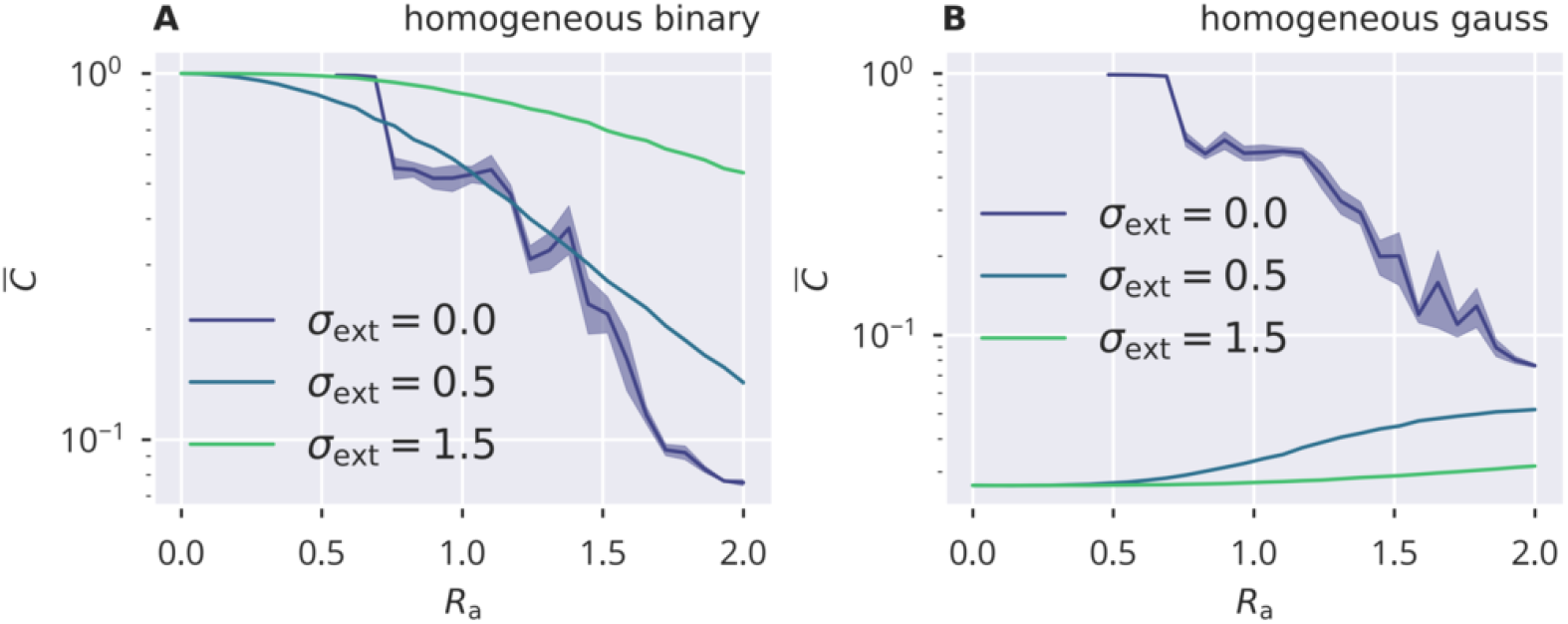
Input induced activity correlations. For homogeneous binary and Gaussian inputs (A/B), the dependency of mean activity cross correlations 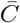, see Eq. (24). 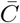 is shown as a function of the target spectral radius *R*_a_. Results are obtained for *N* = 500 sites by averaging over five trials, with shadows indicating the accuracy. Correlations are due to finite-size effect for the autonomous case *σ*_ext_ = 0.

## S1 Appendix. Extended theory of flow control for independent neural activity

We would like to show that the stationarity condition in Eq. (13) results in the correct spectral radius, under the special case of independently identically distributed neural activities with zero mean.

We start with Eq. (13) as a stationarity condition for a given *R*_t_:

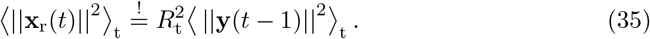

We can express the left side of the equation as

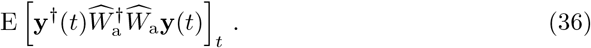

We define 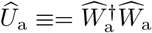 with 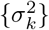 being the set of eigenvalues, which are also the squared singular values of 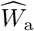, and {**u**_*k*_} the respective set of orthonormal (column) eigenvectors. We insert the identity 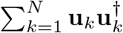 and find

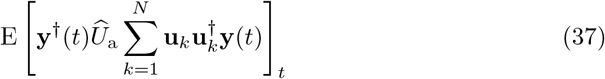

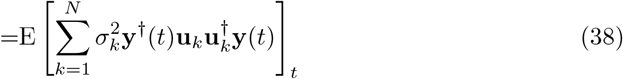

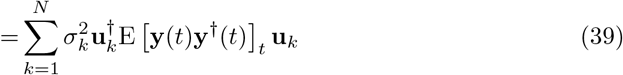

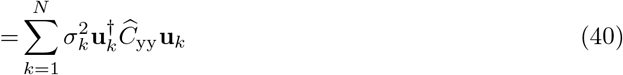

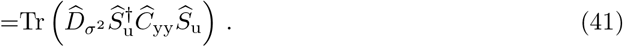

Given zero mean neural activity, 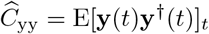 is the covariance matrix of neural activities. 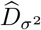 is a diagonal matrix holding the 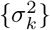 and 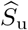 is a unitary matrix whose columns are {**u**_*k*_}. 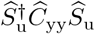 is expressing 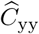 in the diagonal basis of 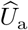.

Including the right hand side of the equation, we get

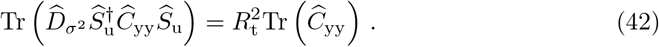

However, since the trace is invariant under a change of basis, we find

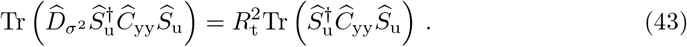

Defining 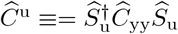, we get

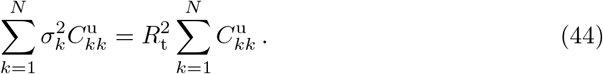

If we assume that the node activities are independently identically distributed with zero mean, we get 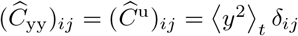. In this case, which was also laid out in Sect., the equation reduces to

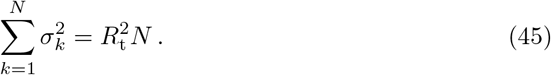

The Frobenius norm of a square Matrix 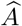 is given by 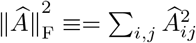. Furthermore, the Frobenius norm is linked to the singular values via 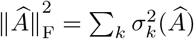 [36, 37]. This allows us to state

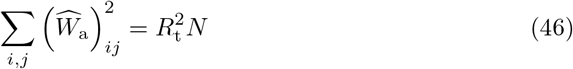

which, by using (11), gives

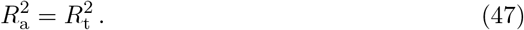

A slightly less restrictive case is that of uncorrelated but inhomogeneous activity, that is 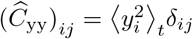. The statistical properties of the diagonal elements 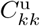 then shows an example of a randomly generated realization of 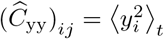 and the resulting diagonal elements of 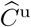, where the corresponding orthonormal basis 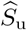 was generated from the SVD of a random Gaussian matrix.

As one can see, the basis transformation has a strong smoothing effect on the diagonal entries, while the mean over the diagonal elements is preserved. Note that this effect was not disturbed by introducing random row-wise multiplications to the random matrix from which the orthonormal basis was derived. The smoothing of the diagonal entries allows us to state that 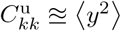 is a very good approximation in the case considered, which therefore reduces (44) to the homogeneous case previously described. We can conclude that the adaptation mechanism also gives the desired spectral radius under uncorrelated inhomogeneous activity.

In the most general case, we can still state that if 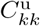 and 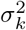 are uncorrelated, for large *N*, Eq. (44) will tend towards

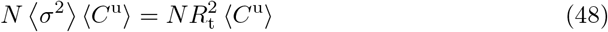

which would also lead to Eq. (45). However, we can not generally guarantee statistical independence since the recurrent contribution on neural activities and the resulting entries of 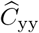 and thus also 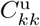 are linked to 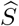 and 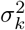, being the SVD of the recurrent weight matrix.

## S2 Appendix Mean field theory for echo state layers

In the following, we deduce analytic expressions allowing to examine the state of echo-state layers subject to a continuous timeline of inputs. Our approach is similar to the one presented by Massar [47].

The recurrent part of the input *x*_*i*_ received by a neuron is a superposition of 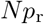 terms, which are assumed here to be uncorrelated. Given this assumption, the self-consistency equations

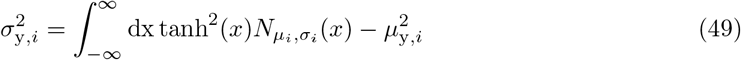

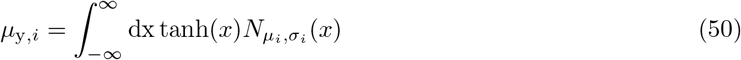

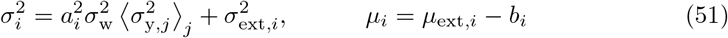

determine the properties of the stationary state. We recall that *σ*_w_ parameterizes the distribution of bare synaptic weights via (26). The general expressions (49) and (50) hold for all neurons, with the site-dependency entering exclusively via *a*_*i*_, *b*_*i*_, *σ*_ext,*i*_ and *μ*_ext,*i*_, as in (51), with the latter characterizing the standard deviation and the mean of the input. Here, 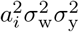 is the variance of the recurrent contribution to the membrane potential, *x*, and *σ*^2^ the respective total variance. The membrane potential is Gaussian distributed, as *N*_*μ,σ*_(*x*), with mean *μ* and variance *σ*^2^, which are both to be determined self-consistently. Variances are additive for stochastically independent processes, which has been assumed in (51) to be the case for recurrent activities and the external inputs. The average value for the mean neural activity is *μ*_*i*_.

For a given set of *a*_*i*_, *σ*_ext,*i*_ and *b*_*i*_, the means and variances of neural activities, 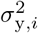 and *μ*_y,*i*_, follow implicitly.

We compared the numerically determined solutions of (49) and (50) against full network simulations using, as throughout this study, *N* = 500, *p*_r_ = 0.1, *σ*_w_ = 1, *μ*_t_ = 0.05. In Fig. 9, the spectral radius *R*_a_ is given for the four input protocols defined in Sect. *Input protocols*. The identical ensemble of input standard deviations *σ*_ext,*i*_ enters both theory and simulations.

Theory and simulations are in good accordance for vanishing input. Here, the reason is that finite activity levels are sustained in an autonomous random neural network when the ongoing dynamics is chaotic and hence decorrelated. For reduced activity levels, viz for small variances 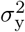, the convergence of the network dynamics is comparatively slow, which leads to a certain discrepancy with the analytic prediction (see Fig. 9).

## Gaussian approximation

The integral occurring in the self-consistency condition (49) can be evaluated explicitly when a tractable approximation to the squared transfer function tanh^2^() is available. A polynomial approximation would capture the leading behavior close to the origin, however without accounting for the fact that tanh^2^() converges to unity for large absolute values of the membrane potential. Alternatively, an approximation incorporating both conditions, the correct second-order scaling for small, and the correct convergence for large arguments, is given by the Gaussian approximation

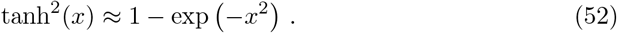

With this approximation the integral in (49) can be evaluated explicitly. The result is

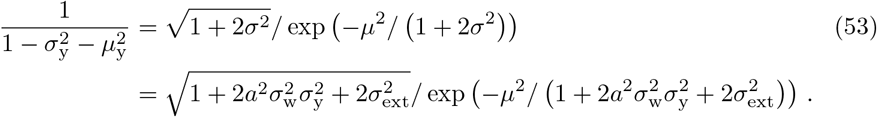

Assuming that *μ* ≈ 0 and *μ*_y_ ≈ 0, inverting the first equation yields a relatively simple analytic approximation for the variance self-consistency equation:

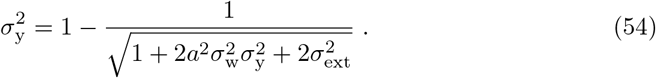

This equation was then used for the approximate update rule in (6) and (18).

